# Mannosylated nanoparticle immunogens enhance the circumsporozoite protein-specific B cell response and improve protection against sporozoite challenge

**DOI:** 10.64898/2026.05.04.722690

**Authors:** Courtney E. McDougal, Mark D. Langowski, Daria Nikolaeva, Chengbo Chen, Derek J. Bangs, Adian Valdez, Rashmi Ravichandran, Suna Cheng, Mariah Lofgren, Marlon Dillon, Delante Sanders, Alvaro Molina-Cruz, Azza H. Idris, Kazutoyo Miura, Kelly Lee, Robert A. Seder, Marion Pepper, Neil P. King

**Author notes:** These authors contributed equally: Courtney E. McDougal, Mark D. Langowski.

## Abstract

Underprocessed oligomannose glycans on protein nanoparticle immunogens engage the innate immune system through mannose-binding lectin and complement, enhancing immunogen trafficking and B cell responses. However, the extent to which oligomannose glycans directly improve protective immunity has remained unclear. Here we generate a series of CSP-bearing I53-50 nanoparticle malaria vaccine candidates with defined numbers and types of engineered N-linked glycans and systematically evaluate their immunogenicity and protective efficacy. Oligomannose display enhanced early plasmablast and germinal center B cell responses, leading to increased CSP-specific memory B cells, long-lived plasma cells, and durable serum antibody titers. Furthermore, nanoparticles bearing oligomannose glycans conferred the strongest protection against sporozoite challenge. By comparing immunogens with defined glycoforms, we attribute improved immune responses and protection specifically to oligomannose rather than complex or truncated glycans. These results will help guide the development of general strategies for glycan engineering aimed at enhancing the protective efficacy of nanoparticle vaccines.

## Introduction

Malaria caused by *Plasmodium falciparum* is a significant public health burden, causing approximately 610,000 deaths in 2024^1^. Use of insecticide-treated bednets and malaria chemoprevention have reduced mortality, but growing drug and insecticide resistance threaten these gains, underscoring the need for a protective vaccine^2^. Two vaccines recently recommended by the WHO, RTS,S/AS01 and R21/Matrix-M, showed efficacies of 55.8% and 68% after 1 year, respectively^3,4^. Both vaccines include a truncated segment of the major surface antigen of the *Plasmodium* sporozoite stage, the circumsporozoite protein (CSP), displaying ∼18 NANP major repeats and the C-terminal domain on Hepatitis B surface antigen virus-like particles^5,6^. These vaccines are currently distributed in areas of moderate-to-high malaria transmission intensity to reduce case incidence and death in children^2^. However, they require three doses to achieve a high anti-CSP antibody titer and the WHO recommends four doses to provide maximum protection in children. Furthermore, because vaccine-elicited antibody titers rapidly wane^4,7–9^, a yearly booster is recommended to maintain efficacy. With time, even boosters are unable to restore antibody titers^10^, highlighting the necessity of developing next-generation malaria vaccines that induce durable, long-lived humoral immunity without requiring continual boosting.

Recently, several studies have demonstrated that the presence of oligomannose glycans on protein nanoparticle immunogens improves humoral responses by enhancing immunogen trafficking to the germinal center (GC)^11–13^. Oligomannose glycans are rare in mature mammalian glycoproteins, whereas this underprocessed glycoform is commonly found in pathogens such as viruses and fungi and is detected as a danger signal by the innate immune system^14^. This occurs through the lectin pathway of complement, where oligomannose-bearing immunogens are opsonized by mannose-binding lectin (MBL), trafficked to lymph nodes, and taken up by subcapsular sinus macrophages. They are then transported to follicular dendritic cells (FDCs), where they are protected from protease activity inside the follicle^15^ and are presented to GC B cells^11,12^. This effect was recently exploited to improve antibody responses against CSP by engineering two oligomannose glycans into a ferritin-based nanoparticle immunogen, 145S^13^. Mice immunized with 145S mounted robust cellular and humoral immune responses and exhibited sterile protection against mosquito bite challenge with transgenic parasites nearly a half year after immunization. However, no head-to-head comparisons were made against nanoparticles bearing different glycoforms, precluding evaluation of the contribution of the oligomannose glycans specifically to vaccine performance. Conversely, we recently showed that engineering N-linked oligomannose glycans—but not complex glycans—onto the surface of a computationally designed protein nanoparticle, I53-50^16^, improved nanoparticle trafficking to B cell follicles in a glycan density- and complement-dependent manner^12^. However, in that study there was no antigen displayed on the glycosylated I53-50 nanoparticles, so we were unable to determine whether improved trafficking influenced the functionality of the resultant immune responses.

Here we prepared and evaluated a series of CSP nanoparticle immunogens with precisely controlled N-linked glycan content and composition. We found that the presence of engineered N-linked glycans improved several aspects of vaccine-elicited humoral immunity—including enhancing germinal center output and protection against sporozoite challenge—and that these improvements were specifically driven by oligomannose glycans.

## Results

### Design and characterization of glycosylated CSP nanoparticles

We hypothesized that the impact of engineered nanoparticle scaffold glycosylation on humoral responses would be most evident when displaying a non-glycosylated antigen, such that added scaffold glycans would not be redundant. We therefore chose to display the repeat region and C terminus of CSP (the “RT” antigen present in RTS,S and R21) on glycosylated I53-50 nanoparticles. I53-50 is a computationally designed, two-component protein nanoparticle comprising 60 trimeric (I53-50A) and 60 pentameric (I53-50B) subunits that assemble into a 120-subunit complex with icosahedral symmetry^16^. We previously generated I53-50 variants with precisely tunable glycan numbers and composition^12^. The most densely glycosylated variant, I53-50A-4gly, contained four N-linked glycans per trimer subunit, for a total of 240 glycans per nanoparticle. We genetically fused RT via a flexible 8-residue Gly-Ser linker (RT.2) to I53-50A-4gly to generate the RT.2-I53-50A-4gly trimeric component (**Fig. 1a**; the amino acid sequences of all novel proteins used in this study are provided in **Supplementary Table 1**). We also produced an RT.2-I53-50A-3gly variant (**Fig. 1a,b**) in which we mutated away the N-linked glycosylation site at position 280—near the nanoparticle interface—that reduced nanoparticle assembly efficiency in our previous study^12^. We further generated a non-glycosylated variant in which all engineered N-linked glycosylation sites were reverted to the original I53-50A sequence, RT.2-I53-50A(No-gly). When assembled, these components would generate I53-50 nanoparticles bearing 60 copies of RT.2 each, as well as 240, 180, and 0 N-linked glycans, respectively (**Fig. 1b,c**).

**Figure 1.**
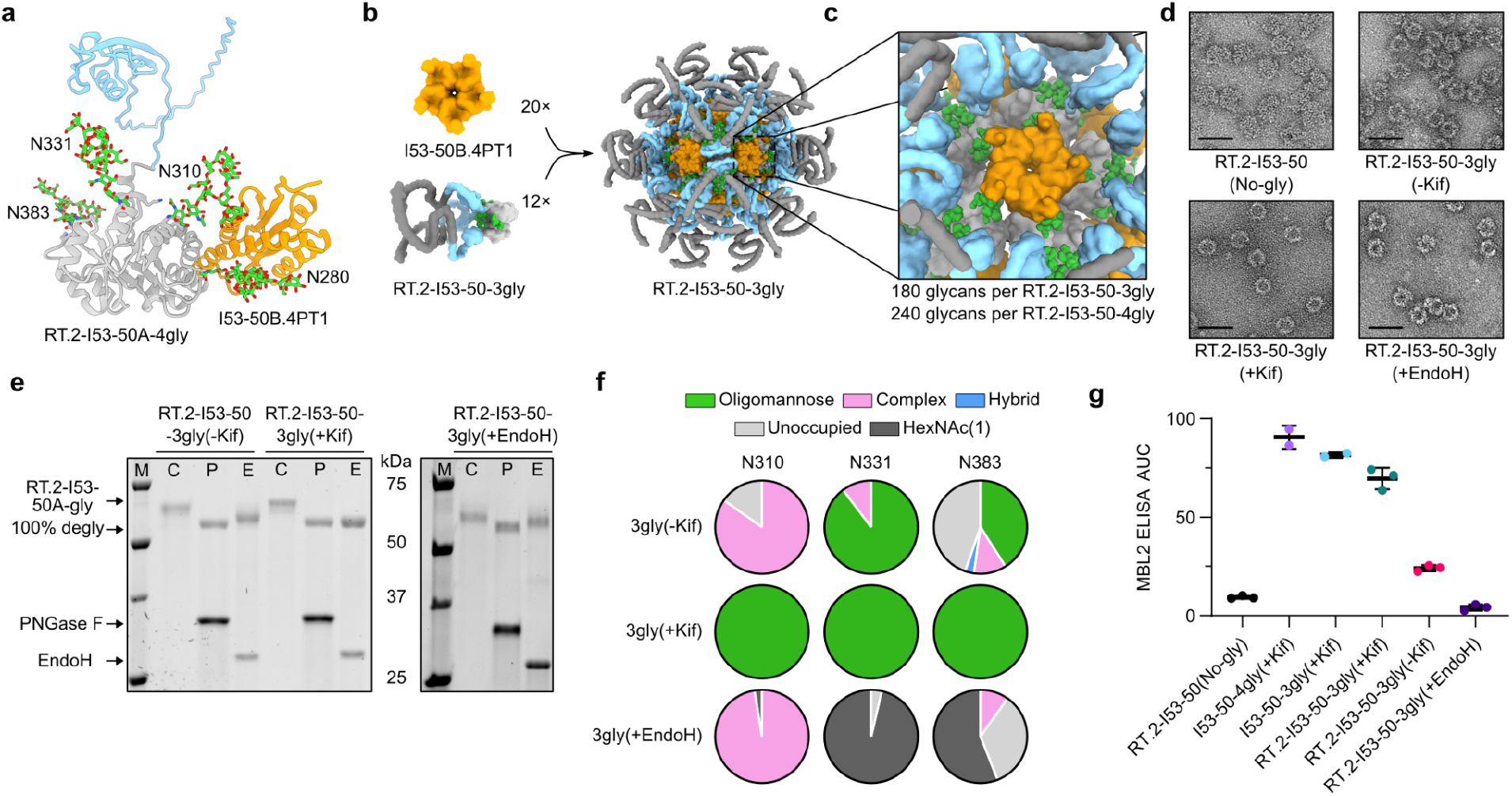
Design and characterization of glycosylated CSP nanoparticles. **a** Model of RT.2-I53-50-4gly asymmetric unit. C terminus of CSP in light blue, I53-50A in light gray, GlcNAc_2_Man_8_ glycans in green, and I53-50B.4PT1 in orange. The NANP repeats of the CSP antigen are not shown for simplicity. **b** AF3 model for RT.2-I53-50A-3gly (major repeats in gray, CSP C terminus light blue, I53-50A in light gray, and glycans in green) and I53-50B.4PT1 (orange). A fully assembled nanoparticle (right) displays 60 copies of the RT antigen. **c** Zoomed-in view of the nanoparticle surface. **d** Negative-stain electron micrographs of nanoparticles at 57k magnification. Scale bars, 50 nm. **e** Glycosidase SDS-PAGE gel shift assay. Control untreated (C), PNGase F (P)-treated, and Endo H (E)-treated glycosylated RT.2-I53-50-3gly nanoparticles. **f** Bottom-up mass spectrometry data for glycosylated RT.2-I53-50A-3gly components at each potential N-linked glycan site. **g** Anti-mouse MBL2 ELISA average (n=3 technical replicates) area under the curve (AUC) measurements against RT.2-I53-50 nanoparticles with and without glycans, and in comparison to non-antigen-bearing I53-50-4gly and -3gly preparations made with kifunensine. RT.2-I53-50-3gly(+EndoH) is a kifunensine-free preparation treated with Endo H to remove all oligomannose glycans. Mean and standard deviation are plotted.

To produce nanoparticles with varying glycan numbers and composition, we first secreted the RT.2-I53-50A-4gly, -3gly, and No-gly components from Expi293F cells in the presence (+Kif) or absence (-Kif) of kifunensine, an inhibitor of ER mannosidases that prevents processing and results in uniformly oligomannose N-linked glycans^17,18^. To generate control nanoparticles that retained complex glycans but were devoid of any oligomannose glycans, we also prepared trimeric components in which we treated purified RT.2-I53-50A-3gly(-Kif) trimers with EndoH, an endoglycosidase that specifically cleaves oligomannose glycans to leave behind only a HexNAc(1) core. We purified each protein by immobilized metal affinity chromatography (IMAC) and size exclusion chromatography (SEC), and then assembled them into I53-50 nanoparticles by mixing with I53-50B.4PT1 pentamer in a 1.1:1 molar ratio (**Fig. 1b**). After the assembled nanoparticles were separated from excess components by SEC, SDS-PAGE confirmed that the preparations contained both of the expected bands and were highly pure (**Supplementary Fig. 1a,b**). The presence of assembled nanoparticles with diameters ranging from 33-38 nm was further confirmed by dynamic light scattering (DLS) and negative-stain electron microscopy (nsEM) (**Fig. 1d & Supplementary Fig. 1c,d**). Not unexpectedly, the RT.2-I53-50-4gly nanoparticles assembled less efficiently than the others, as evidenced by considerable trailing shoulders after the nanoparticle peak and relatively large component peaks in their SEC chromatograms. As a result, we focused further studies on the RT.2-I53-50-3gly variants.

### Glycoprofiling of glycosylated CSP nanoparticles

We next estimated the glycan composition of our nanoparticles by glycosidase gel shifts during SDS-PAGE. We treated each SEC-purified nanoparticle with PNGase F to eliminate all N-linked glycans or EndoH to reduce oligomannose glycans to a single HexNAc(1) while leaving complex glycans intact. In every case, PNGase F treatment resulted in a single RT.2-I53-50A band that migrated more rapidly than untreated samples as expected (**Fig. 1e** and **Supplementary Fig. 1e**). The same effect was observed when the (+Kif) nanoparticles were treated with EndoH, indicating they contained uniformly oligomannose glycans. By contrast, treatment of the (-Kif) nanoparticles with EndoH caused only a partial shift, suggesting that they contained some oligomannose and some complex glycans.

Analysis of N-linked glycosylation occupancy by mass spectrometry confirmed that 3gly(+Kif) nanoparticles had 100% oligomannose at all N-linked glycan positions as expected (**Fig. 1f**). In the RT.2-I53-50-3gly(-Kif) nanoparticles, the glycans at position 310 were >75% complex with no oligomannose detected, but positions 331 and 383 contained 89% and 41% oligomannose, respectively, explaining the partial gel shift observed upon EndoH treatment (**Fig. 1e,f**). The high levels of oligomannose observed at these positions differed from our previous glycoprofiling of I53-50-4gly without displayed antigen, which revealed >75% complex glycans at all sites when prepared without kifunensine^12^. These results suggest that N-terminal display of the RT antigen may sterically occlude mannosidase enzymes from processing the glycans in I53-50A, particularly at the nearby position 331 (**Fig. 1a**), resulting in the presence of underprocessed oligomannose glycans in the secreted glycoprotein. A similar phenomenon has been observed in many densely glycosylated viral glycoproteins^19,20^.

Lastly, we measured the binding of murine mannose-binding lectin 2 (MBL2) to each nanoparticle using an MBL2 enzyme-linked immunosorbent assay (ELISA) to gauge whether they may engage MBL and the complement pathway *in vivo*. We found that the level of MBL2 binding to each nanoparticle correlated with its oligomannose content: the +Kif nanoparticles exhibited the highest binding, RT.2-I53-50-3gly(-Kif) showed some residual binding, and RT.2-I53-50-3gly(+EndoH) did not bind MBL2, as expected (**Fig. 1g**). Together, these data demonstrate the preparation of a series of antigen-bearing nanoparticle immunogens with precisely varying glycan number and composition.

### Mannosylation of I53-50 nanoparticles improves acute B cell numbers and differentiation

To begin investigating the impact of scaffold mannosylation on B cell responses to vaccination, we measured acute B cell responses in mice receiving 1) the unglycosylated RT.2-I53-50(No-gly), 2) the glycosylated but mannose-deficient RT.2-I53-50-3gly(+EndoH), and 3) the mannosylated RT.2-I53-50-3gly(+Kif) nanoparticles. We compared responses to these immunogens in C57BL/6 mice and mice deficient in the essential complement component C3 (C3KO), as previous studies have indicated that oligomannose glycoforms enhance B cell responses via complement^11,12^. We vaccinated mice intramuscularly (IM) with 3 µg of each nanoparticle formulated in SMNP adjuvant^21^ and harvested draining lymph nodes (dLN) 8 days later (**Fig. 2a**). Using CSP-containing B cell tetramers, we found that C57BL/6 mice immunized with 3gly(+Kif) nanoparticles showed an increased number of CSP^+^ B cells compared to mice immunized with the No-gly and 3gly(+EndoH) nanoparticles (**Fig. 2b** and **Supplementary Data Fig. 2**). This effect was lost in C3KO mice, confirming the previous finding that oligomannose glycans enhance acute antigen-specific B cell numbers in a complement-dependent manner^11,12^ (**Fig. 2b**).

**Figure 2.**
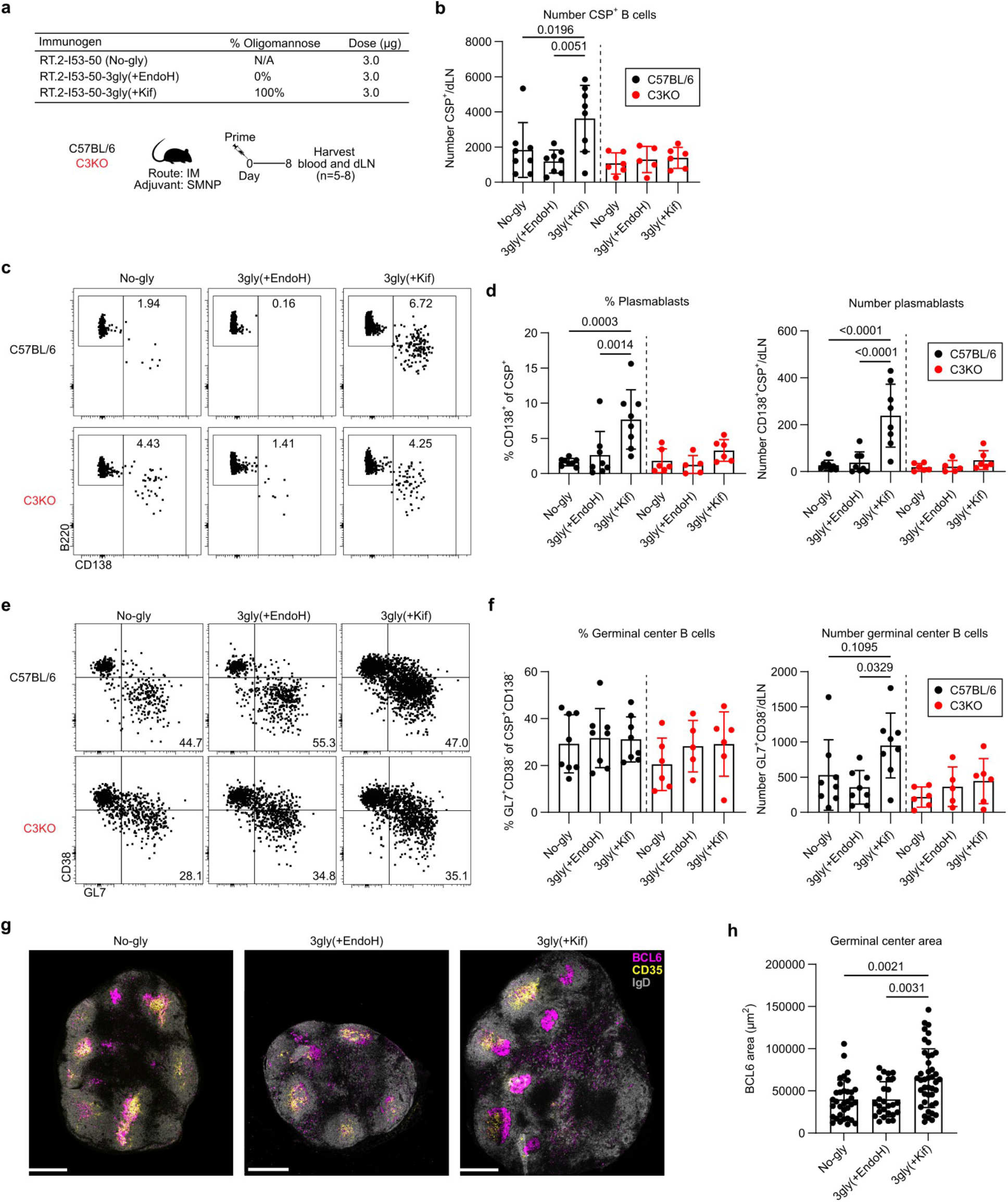
Mannosylation of I53-50 nanoparticles improves acute B cell numbers and differentiation. **a** C57BL/6 or C3-deficient (C3KO) mice were immunized intramuscularly with 3 µg of No-gly, 3gly(+EndoH), or 3gly(+Kif) nanoparticles with SMNP adjuvant. Eight days later, blood and inguinal lymph nodes (dLn) were harvested for analysis. n=5-8. **b** The total number of CSP^+^ B cells isolated from the dLn. **c** Representative plots, **d** frequency (left), and total number (right) of CSP-specific PBs (CSP^+^CD138^+^) isolated from the dLn. **e** Representative plots, **f** frequency (left), and total number (right) of CSP-specific GC B cells (CSP^+^CD138^−^CD38^−^GL7^+^) isolated from the dLn. **g** Representative staining to identify follicular dendritic cells (CD35^+^), GC B cells (BCL6^+^), and naive B cells (IgD^+^). Scale bar, 500 µm. **h** Quantification of the GC area (BCL6^+^). n=3 mice, 3 images per dLN. Statistical analysis by 2-way ANOVA with a Benjamini-Hochberg correction. Mean and standard deviation are plotted.

We next wanted to understand if the addition of oligomannose glycans on the nanoparticles would alter the differentiation of CSP-specific B cells in addition to their number. We found that eight days after immunization, C57BL/6 mice vaccinated with 3gly(+Kif) nanoparticles generated a significantly higher number and proportion of CSP^+^CD138^+^ plasmablasts (PBs) compared to mice receiving the No-gly and 3gly(+EndoH) nanoparticles (**Fig. 2c,d**). This phenotype was not observed in C3KO mice, again demonstrating dependence on complement (**Fig. 2c,d**).

Because we detected elevated levels of CSP^+^ PBs after vaccination with 3gly(+Kif) nanoparticles in C57BL/6 mice, we next looked in the serum to see if there was a concurrent increase in CSP-specific antibody titers. We found that mice vaccinated with 3gly(+Kif) had significantly higher levels of CSP-specific serum IgG compared to those vaccinated with 3gly(+EndoH) (**Supplementary Fig. 3a,b**). Vaccination of C3KO mice generated very little antibody response (**Supplementary Fig. 3a,b**), consistent with the observed dependence of the PB response on complement (**Fig. 2c,d**).

We next examined if CSP-specific B cells entered germinal centers (GCs). Though there was no difference in the frequency of CSP^+^CD38^−^GL7^+^ GC B cells among CD138^−^ (non-PB) CSP^+^ B cells, we observed an increase in the total number of CSP^+^ GC B cells in 3gly(+Kif)-vaccinated mice compared to those receiving the No-gly or 3gly(+EndoH) nanoparticles (**Fig. 2e,f**). This phenotype was highly dependent on complement, as vaccination of C3KO mice with mannosylated nanoparticles did not induce an enhanced GC response (**Fig 2e,f**). Consistently, when we analyzed the size of GCs (defined as BCL6^+^) by confocal microscopy, we found that 3gly(+Kif)-vaccinated C57BL/6 mice had significantly larger GCs compared to those receiving No-gly or 3gly(+EndoH) nanoparticles (**Fig. 2g,h**).

To assess the generality of the improved B cell response to mannosylated nanoparticle immunogens, we replicated our study using a different route of administration, adjuvant, and dose. We vaccinated C57BL/6 mice with 18 µg No-gly or 3gly(+Kif) nanoparticles formulated in Sigma Adjuvant System (SAS) via intraperitoneal (IP) injection (**Supplementary Data Fig. 4a**). Eight days later, we harvested spleens and found that, consistent with our prior experiments, mannosylated immunogens enhanced the total number of vaccine-elicited CSP^+^ B cells and CSP-specific PBs and GC B cells (**Supplementary Data Fig. 4b,c,e**). We also detected increased titers of CSP-specific serum IgG by ELISA (**Supplementary Data Fig. 4d**). In sum, our data show that mannosylated RT.2-I53-50 nanoparticles induced increased numbers of CSP-specific antibody-secreting PBs and GC B cells compared to non-mannosylated nanoparticles, and that this effect is consistent across different vaccination regimens.

### Mannosylated nanoparticles generate elevated levels of CSP-specific memory B cells, long-lived plasma cells, and durable antibody titers

Having established that display of oligomannose glycans on RT.2-I53-50 improves acute B cell responses, we next assessed its effect on the establishment of long-lived humoral responses. We vaccinated C57BL/6 and C3KO mice IM with 3 µg of No-gly, 3gly(+EndoH), or 3gly(+Kif) nanoparticles formulated in SMNP and harvested spleens, draining lymph nodes, and blood at least 60 days later (60-65 days, experiment dependent) (**Fig. 3a**).

We first assessed the memory B cell (MBC) compartment. The presence of oligomannose glycans did not significantly alter the number of class-switched CSP-specific B cells (**Fig. 3b,c**). However, we found that the 3gly(+Kif) nanoparticles generated significantly elevated numbers of CD73^+^CD80^+^ CSP-specific MBCs compared to No-gly and 3gly(+EndoH) particles (**Fig. 3d,e**). These markers define highly functional MBCs poised to secrete antibody rapidly upon restimulation^22–26^. This effect was again complement-dependent, as the elevated CD73^+^CD80^+^ population induced by the 3gly(+Kif) nanoparticles was lost in C3KO mice (**Fig. 3d,e**). These data suggest that addition of mannose improves CSP-specific MBC responses in a complement-dependent manner.

**Figure 3.**
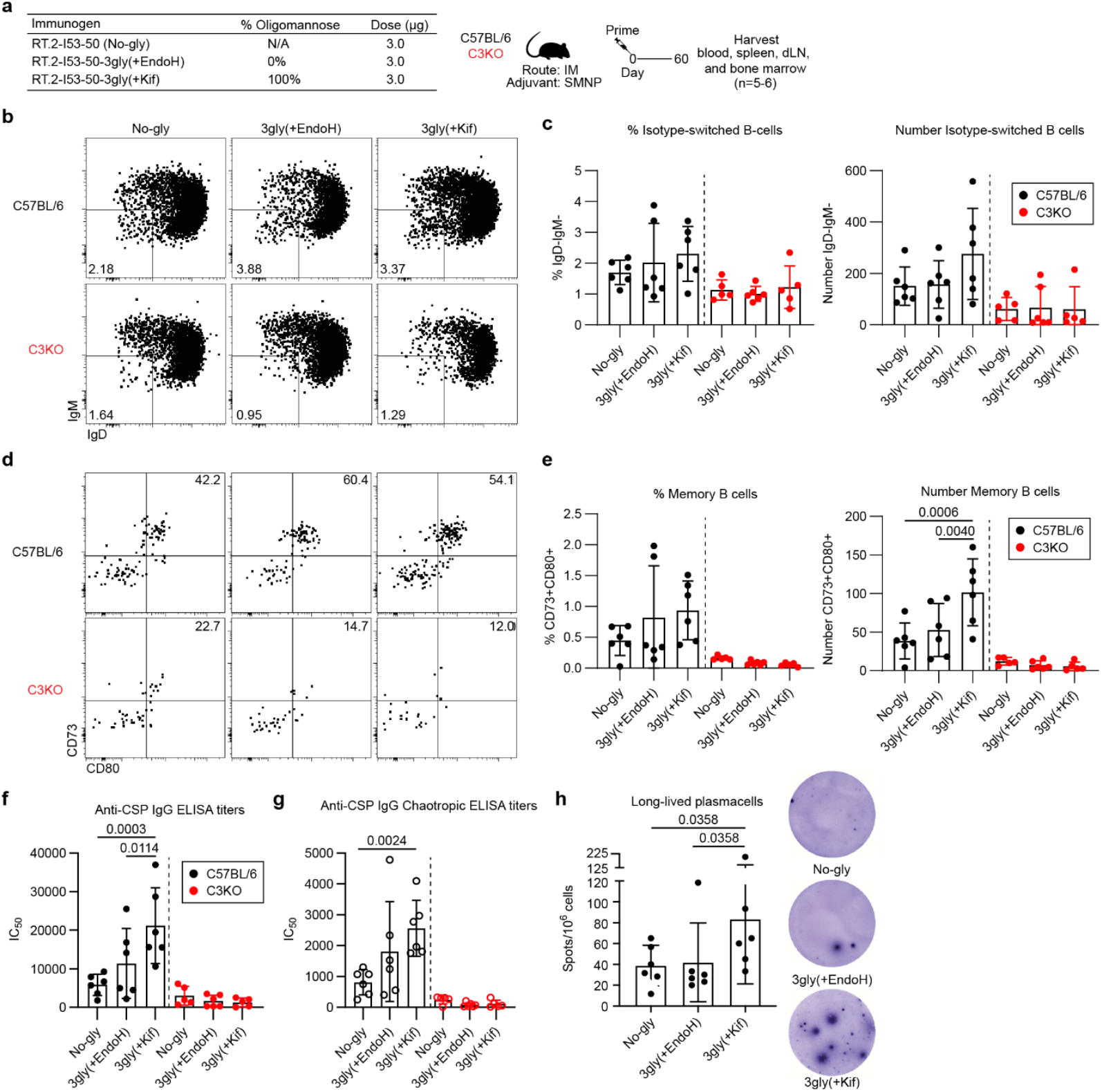
Mannosylation of I53-50 nanoparticles improves MBC and LLPC responses. **a** C57BL/6 or C3-deficient (C3KO) mice were immunized intramuscularly with 3 µg of No-gly, 3gly(+EndoH), or 3gly(+Kif) nanoparticles with SMNP adjuvant. 60-65 days later, blood was harvested for serum antibody analysis, inguinal lymph nodes (dLn) and spleens were pooled to isolate CSP-specific memory B cells, and bone marrow was harvested to detect LLPCs. **b** Representative plots and, **c** quantification of the proportion (% of CSP^+^CD138^−^GL7^−^CD38^+^ B cells) and total number of CSP-specific class-switched (IgM^−^IgD^−^) B cells in the spleen and dLn. **d** Representative plots and, **e** quantification of the proportion (% of CSP^+^CD138^−^GL7^−^CD38^+^ B cells) and total number of CSP-specific CD73^+^CD80^+^ B cells in the spleen and dLn. **f** Serum CSP-specific IgG was quantified by ELISA. **g** High-affinity serum CSP-specific IgG by chaotropic ELISA using a urea wash. **h** CSP-specific IgG LLPCs in the bone marrow were detected by ELISpot. n=5-6. Statistical analysis by 2-way ANOVA with a Benjamini-Hochberg correction. Mean and standard deviation are plotted.

Sustained serum antibody titers are particularly important for preventing *Plasmodium* infection, where sporozoites can invade hepatocytes within hours, limiting the time available for MBC recall. Accordingly, we next looked 60 days after immunization for maintained CSP-specific serum antibody by ELISA and found that immunization of C57BL/6 mice with 3gly(+Kif) resulted in significantly higher CSP-specific IgG titers compared to the non-mannosylated immunogens in a complement-dependent manner (**Fig. 3f**). In addition to increased total antigen-specific IgG, 3gly(+Kif) nanoparticle vaccination resulted in elevated levels of high-affinity antibodies compared to No-gly particles as measured by chaotropic ELISA (**Fig. 3g**), which uses a urea wash to dislodge low-affinity antibodies. This suggests that the increased GC B cell responses we observed earlier (**Fig. 2e-h**) resulted in productive affinity maturation. Complement deficiency again reduced the high-affinity titers (**Fig. 3g**).

In the absence of an active infection, long-lived plasma cells (LLPCs) are responsible for maintaining levels of circulating antibodies. As high antibody titers in 3gly(+Kif) immunized mice were maintained well after the initial PB burst, we suspected that LLPCs were likely the source of the durable antibody titers. To assess LLPC number, we harvested bone marrow from the C57BL/6 mice vaccinated with the three immunogens and quantified the number of LLPCs by ELISpot. Strikingly, 3gly(+Kif) nanoparticles induced a significantly higher number of LLPCs compared to mice immunized with the No-gly or 3gly(+EndoH) particles (**Fig. 3h**). This suggests that mannosylated nanoparticles drive enhanced LLPC formation, leading to elevated antibody titers that may provide superior protection from *Plasmodium* infection. Together, these findings demonstrate that oligomannose display enhances complement-dependent establishment of long-lived humoral immunity by improving GC output including MBC number, LLPC formation, and sustained high-affinity serum antibody responses.

### Mannosylated nanoparticles confer superior protection against parasite challenge

To assess the impact of oligomannose glycans on the functionality of the vaccine-elicited antibody response, we vaccinated groups of C57BL/6 mice IM with a series of nanoparticle immunogens and challenged them with sporozoites (**Fig. 4a**). In addition to the No-gly, 3gly(+EndoH), and 3gly(+Kif) RT.2-I53-50 nanoparticles, we also vaccinated a group of mice with the 3gly(-Kif) nanoparticles produced with neither kifunensine nor Endoglycosidase H treatment (**Fig. 4a**). Though these nanoparticles contained lower levels of oligomannose compared to the 3gly(+Kif) immunogens (**Fig. 1f,g**), they would be more straightforward to manufacture at scale, and we therefore wanted to understand whether their partial oligomannose content would suffice to induce the robust humoral responses seen upon 3gly(+Kif) vaccination. Based on prior characterization of the dynamic range of the sporozoite challenge model used^27^, we primed each group of mice with 3 µg of nanoparticle immunogen formulated in SMNP and boosted them 3 weeks later (**Fig. 4a**).

**Figure 4.**
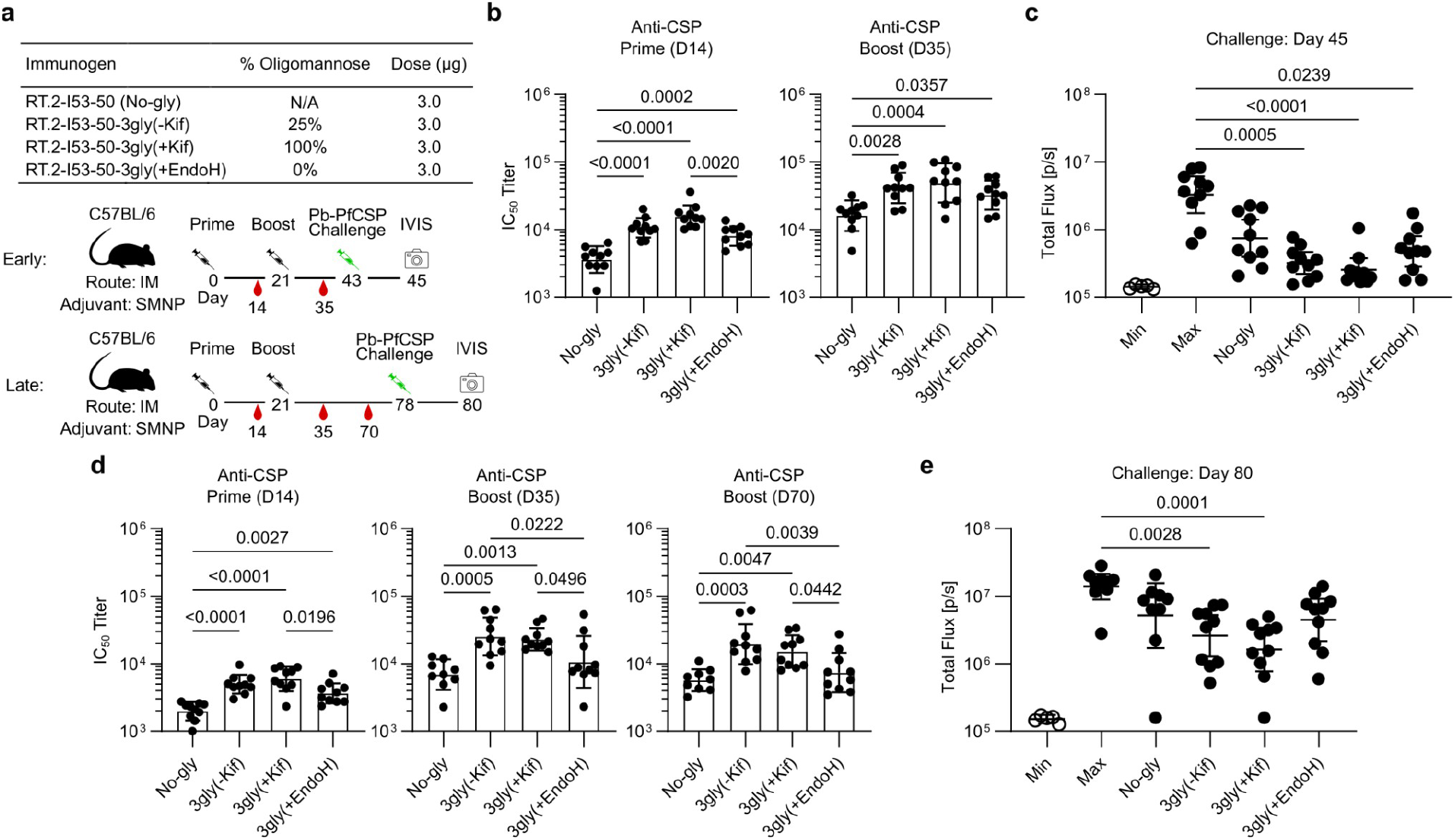
Mannosylated nanoparticles confer superior protection against parasite challenge. **a** Two study arms (early and late challenge) of C57BL/6 mice were immunized intramuscularly with 3 µg of No-gly, 3gly(-Kif), 3gly(+Kif), or 3gly(+EndoH) nanoparticles with SMNP adjuvant and boosted 21 days later. Mice were challenged with 2,000 *P. berghei*-PfCSP-GFP/Luc sporozoites intravenously and liver burden was assessed via IVIS two days later (day 45 for the early group, and day 78 for the late group). **b** Early study serum IgG titers against PfCSP antigen after prime and boost immunizations at days 14 and 35. Statistical analysis by one-way ANOVA with multiple comparisons. **c** Early study total flux in uninfected mice (min), unvaccinated infected mice (max), and mice vaccinated with the indicated nanoparticles. n=5-10. Statistical analysis by Kruskal-Wallis test with multiple comparisons comparing all groups except for the uninfected control. **d** Late study serum IgG titers against PfCSP antigen after prime and boost immunizations at days 14, 35, and 70. Statistical analysis by one-way ANOVA with multiple comparisons. **e** Late study total flux in uninfected mice (min), unvaccinated mice (max), and mice vaccinated with the indicated nanoparticles. n=5-10. Statistical analysis by Kruskal-Wallis test with multiple comparisons comparing all groups except for the uninfected control.

We harvested blood from the mice two weeks after the primary vaccination and boost and assessed anti-CSP IgG antibody titers by ELISA. Consistent with our earlier analyses, we found that the 3gly(+Kif) nanoparticles induced significantly higher levels of anti-CSP antibody than the No-gly nanoparticles at both timepoints, and higher than the glycosylated but mannose-deficient (3gly(+EndoH)) nanoparticles post-prime (**Fig. 4b**). Notably, the 3gly(-Kif) and 3gly(+EndoH) nanoparticles also elicited higher antibody titers than the No-gly nanoparticles at both time points.

To assess the protection conferred by each immunogen, we challenged the mice 22 days post-boost (D45) with 2,000 transgenic *P. berghei*-PfCSP-GFP/Luc sporozoites delivered intravenously (IV), and two days later measured parasite burden in the liver using an *in vivo* imaging system (IVIS)^28^. All three groups of mice that received glycosylated immunogens—but not the mice receiving No-gly nanoparticles—had significantly reduced liver burden compared to unvaccinated control mice (max), with the mannosylated immunogens 3gly(+Kif) and 3gly(-Kif)) showing the largest reductions (**Fig. 4c**).

A major limitation of current malaria vaccines is the limited durability of the humoral responses they elicit^10,29^. Accordingly, we repeated our challenge study with a longer interval between boost and challenge (57 days) to test the effect of oligomannose display on the durability of the response (**Fig. 4a**). Similar trends for anti-CSP IgG titers were observed, except that the mannosylated nanoparticles (3gly(+Kif) and 3gly(-Kif)) elicited higher titers than both the No-gly and 3gly(+EndoH) groups two weeks (D35) and seven weeks (D70) after boosting (**Fig. 4d**).

Liver burden was again measured two days after challenge by IVIS and, strikingly, only the mice receiving mannosylated nanoparticles had significant liver burden reduction compared to unvaccinated mice (**Fig. 4e**). Together, our challenge studies confirmed our earlier observations that mannosylated nanoparticle immunogens elicit superior humoral responses, and showed that these superior responses conferred improved protection from challenge, even 8 weeks later.

## Discussion

Here we show that engineered oligomannose glycans on the surface of CSP-bearing I53-50 nanoparticles enhance vaccine-elicited B cell responses and improve protection against sporozoite challenge. Oligomannose display increased acute plasmablast and germinal center B cell responses, leading to elevated CSP-specific memory B cells, long-lived plasma cells, and serum antibody titers. These effects could be attributed specifically to oligomannose glycans rather than complex or truncated glycoforms. Our findings extend pioneering studies from the Irvine group demonstrating enhanced trafficking of mannosylated nanoparticle immunogens^11,12,30^ by showing that this advantage translates into improved protection against sporozoite challenge. Although a previous study reported improved protection from a CSP-bearing ferritin nanoparticle carrying engineered oligomannose glycans relative to a non-glycosylated comparator^13^, that work did not compare distinct glycoforms, leaving unclear the extent to which oligomannose itself drove the observed effect. By systematically comparing nanoparticle immunogens bearing different numbers and types of N-linked glycans, our study links innate immune recognition of engineered oligomannose glycans directly to improved functional outcomes after vaccination.

Unexpectedly, RT.2-I53-50-3gly nanoparticles produced without kifunensine nevertheless contained substantial oligomannose. In contrast, a similar I53-50A-4gly variant expressed previously without fused antigen carried predominantly complex glycans^12^. We therefore infer that the large, flexible RT antigen sterically restricted glycan-processing enzymes in the secretory pathway, particularly at the N331 site, resulting in increased underprocessed oligomannose^19^. Steric effects also likely influenced MBL engagement. RT.2-I53-50A-3gly produced with kifunensine, which was fully mannosylated at all three sites, bound recombinant MBL strongly, whereas protein produced without kifunensine, in which oligomannose was concentrated primarily at N331, showed reduced binding. Nevertheless, even this partial engagement enhanced immunogenicity: enzymatic removal of oligomannose from RT.2-I53-50-3gly(-Kif) nanoparticles abolished MBL binding and significantly reduced anti-CSP antibody titers. These results demonstrate that appropriately positioned engineered glycans can remain underprocessed and confer the immunogenicity benefits of MBL engagement without kifunensine treatment^31^ or specialized expression systems^32^. This strategy could simplify industrial manufacturing of glycosylated vaccines by enabling the use of standard expression platforms such as CHO cells while retaining the advantages of oligomannose display. Genetically encoded oligomannose glycans may similarly enhance nucleic-acid–delivered immunogens; for example, antigen-derived oligomannose glycans likely contributed to the immunogenicity of eOD-GT8 60mer delivered by mRNA in clinical trials^33^.

Looking forward, the dual advantages of improved immune responses and simplified manufacturing should motivate further efforts to engineer oligomannose glycans into designed immunogens. Our results, together with previous studies, suggest several guidelines for these efforts. First, as asserted previously^13^, engineered oligomannose glycans are likely to be most beneficial for nanoparticle immunogens displaying non-glycosylated antigens, such as those derived from parasites, bacteria, or non-enveloped viruses. This conclusion is supported by both the present work and earlier studies demonstrating improved B cell responses and protection against *Plasmodium* sporozoite challenge^13^, as well as by work from the Irvine laboratory showing that removal of glycans from viral glycoprotein nanoparticles by EndoH or PNGase F treatment impaired trafficking and reduced immunogenicity^11,12^. Second, engineering glycans into nanoparticle scaffold subunits rather than the displayed antigen provides a modular strategy for enhancing antigen-specific responses while minimizing the risk of perturbing antigen structure or antigenicity. This approach should be particularly powerful when combined with methods for de novo generation of nanoparticle scaffolds^34,35^. Third, glycan placement is critical. The requirement for the RT antigen to promote underprocessing at N331—and the reduced MBL binding likely caused by steric hindrance from the displayed antigen—illustrate how glycan location influences accessibility and function. Incorrect placement can also interfere with assembly, as observed for the RT.2-I53-50-4gly nanoparticles. Conversely, we previously showed that an engineered glycan within a designed nanoparticle was produced exclusively as oligomannose due to steric constraints imposed by nanoparticle assembly^34^, suggesting another strategy for deliberately encoding oligomannose in future designs. Recent advances in machine learning-based structural modeling now make it possible to accurately represent proteins together with post-translational modifications such as glycans^36–38^. These methods should enable more precise engineering of oligomannose glycans and facilitate the design of nanoparticle vaccines that optimally engage the innate immune system while avoiding unintended effects on assembly or stability.

There are several limitations of our study that suggest important directions for future research. First, while murine models enable initial evaluation of new vaccine technologies and candidates in a fully functioning immune system, it remains essential to validate these findings in humans. Encouragingly, a previous study demonstrated that mannosylated nanoparticle immunogens efficiently accumulated in B cell follicles following vaccination in non-human primates^30^. Clinical evaluation will therefore be an important next step for validating and de-risking this vaccine design strategy. Additionally, as the physiological route of *Plasmodium* infection is via mosquito bite, evaluating protection from mosquito bite challenge would provide relevant insights into vaccine efficacy.

In summary, incorporation of oligomannose glycans on the surface of RT.2-I53-50 nanoparticles enhanced B cell responses and significantly improved protection against malaria in a mouse model. Our findings highlight the potential of oligomannose-bearing nanoparticles as a promising platform for the development of next-generation malaria vaccines with enhanced immunogenicity and durable protection. More broadly, our results showcase how engineered glycans can be used to engage the innate immune system and define principles to guide future efforts in glycoengineering nanoparticle vaccines.

## Methods

### Molecular modeling

The structures of the RT.2-I53-50A-3gly and -4gly nanoparticle components were predicted with AlphaFold3^36^ with GlcNAc_2_Man_6_ glycans at each N-linked glycan sequon. The full RT.2-I53-50-3gly nanoparticle was modeled in icosahedral symmetry with ChimeraX 1.6 software (UCSF) using the RT.2-I53-50A-3gly AF3 prediction in combination with a structure of I53-50B.4PT1 (PDB ID: 6P6F).

### Plasmid synthesis

Plasmids for I53-50B.4PT1 and RT-I53-50A were constructed using the pET29b+ vector as previously described^39^. For mammalian-expressed non-glycosylated and glycosylated components, the RT antigen was genetically fused by an 8-residue GS linker (RT.2) to I53-50A or I53-50A-4gly (RT.2-I53-50A or RT.2-I53-50-4gly, respectively; **Supplementary Table 1**), the latter of which was based off of a previously described construct with four N-linked glycan sequons^12^. RT.2-I53-50-3gly variants were produced by mutating the first glycan sequon to the original I53-50A sequence^16^. Constructs were codon-optimized for mammalian expression with a bovine prolactin signal sequence and a C-terminal hexahistidine tag, and inserted into a pCMV/R vector (XbaI and AvrII restriction sites) by Genscript.

### Nanoparticle protein expression

I53-50B.4PT1 and RT-I53-50A components were expressed as previously described in *E. coli*^*16,39*^. RT.2-I53-50A and glycosylated RT.2-I53-50A variant plasmids were used to transiently transfect Expi293F cells using PEI-MAX (Polysciences). Glycosylated RT.2-I53-50A variants were expressed without (referred to as standard or kifuensine-free preparations; designated “-Kif”) or with 5 µM kifunensine, the latter of which results in 100% oligomannose glycoforms (designated “+Kif”). Kifunensine-treated preparations were supplemented with 25 mM glucose to improve N-linked glycan occupancy.

### RT.2-I53-50A component purification

After expression, mammalian cultures were centrifuged and supernatant was collected for downstream purification. Ni Sepharose Excel resin was added (10 mL per 200 mL supernatant) to each supernatant, and flowthrough was collected via a gravity column. Resin was subsequently washed with 10 column volumes (CV) of 50 mM Tris, 500 mM NaCl, 20 mM Imidazole, 0.75% CHAPS, pH 8.0 (wash buffer). Bound components were eluted with 50 mM Tris, 500 mM NaCl, 300 mM Imidazole, 0.75% CHAPS, pH 8.0 (elution buffer) in two 2.5 CV fractions, the first of which contained the protein of interest. The eluted components were further purified by SEC using a Superdex 200 Increase 10/300 GL column (Cytiva) in 25 mM Tris, 150 mM NaCl, 0.75% CHAPS, pH 8.0 (component sizing buffer). 1 mL fractions were collected for peaks that corresponded to the expected elution volumes of the components, and their purity was determined by SDS-PAGE. Component concentrations were determined by UV-vis (Agilent Cary 3500), and low endotoxin content confirmed by Endosafe Endotoxin Testing Cartridges (Charles River).

### RT.2-I53-50 nanoparticle assembly and purification

Purified RT.2-I53-50A (bare and glycosylated) components were assembled *in vitro* in a 1.1:1 molar ratio with I53-50B.4PT1 in 25 mM Tris, 150 mM NaCl, 5% Glycerol pH 8.0 (assembly buffer) at a total molar concentration of 20-50 µM as previously described^39^, and left overnight at 4°C on a rocking shaker. Nanoparticles were purified from excess component by SEC using a Superose 6 Increase 10/300 GL column (Cytiva) in assembly buffer. Fractions corresponding to the expected elution volume of fully formed assemblies were collected for downstream analysis. Purity was determined by SDS-PAGE and concentration by UV-vis (Agilent Cary 3500).

### Dynamic light scattering (DLS)

Measurements for nanoparticles were performed on an UNcle Nano-DSF (UNchained Laboratories) using quartz capillary cassettes (UNi, UNchained Laboratories). Four acquisitions were collected using auto attenuation of the laser, and the UNcle analysis software (UNchained Laboratories) was used to determine the hydrodynamic diameter and polydispersity index (PDI) of assemblies.

### Negative-stain electron microscopy (nsEM)

Assembled nanoparticles at a concentration of 50 µg/mL were added to carbon-covered 400 mesh copper grids (Electron Microscopy Sciences) and stained with 2% uranyl formate. Micrographs were imaged on a Talos 120C microscope with a Ceta camera.

### Glycan profiling by glycosidase gel shifts

After denaturation, the N-linked glycan content of glycosylated RT.2-I53-50A components or nanoparticles was estimated by treatment with PNGase F (NEB) which cleaves all N-linked glycoforms, and Endo H (EndoH, NEB) which only cleaves oligomannose. These samples were analyzed by SDS-PAGE and their gel shifts after treatment were compared to an untreated control. A shift in molecular weight after treatment with Endo H indicated the presence of oligomannose glycans.

### Glycan profiling by mass spectrometry (MS)

N-linked glycosylation profiles at each glycosylated RT.2-I53-50A potential N-linked glycan sequon was determined by MS as previously described^12^.

### Murine mannose-binding lectin 2 (MBL2) enzyme-linked immunosorbent assay (ELISA)

Nunc MaxiSorp ELISA plates (ThermoFisher) were coated with 50 µL of 1 µg/mL RT.2-I53-50 or glycosylated RT.2-I53-50 nanoparticles in phosphate-buffered saline (PBS) in triplicate and left covered overnight in a cold room. The next day, the plates were washed 3× with PBS + 0.05% Tween-20 (PBS-T). Recombinant MBL2 (Bio-Techne Corporation) was diluted in D-PBS with calcium and magnesium + 1% Bovine Serum Albumin (blocking buffer), diluted 1:2 down the column of the plate, and incubated for 1 hour at 37°C. After incubation, plates were washed 3× with PBS-T, and anti-MBL2 antibody (14D12, Abcam) in blocking buffer at a concentration of 2 µg/mL was added to each well and incubated at room temperature for 1 hour. Plates were washed 4× with PBS-T, and secondary antibody HRP conjugate (anti-rat IgG:HRP, BioRad) at a 1:6000 dilution in blocking buffer was added to each well and incubated for 1 hour at room temperature. 50 µL of TMB substrate was added to each well and developed for 5 minutes, and subsequently neutralized with sulfuric acid stop solution. Absorbance at 450 nm was measured using an Epoch Microplate reader (Biotek) and values plotted and Area Under Curve (AUC) calculated using GraphPad Prism (GraphPad Software).

### Anti-CSP serum IgG ELISA

Corning 96-well plates were coated overnight at 4°C with 2 μg recombinant SAmut CSP protein^40^ diluted in PBS. Plates were washed with PBST and blocked for one hour at room temperature in PBST + 3% milk. Plasma from murine samples were serial-diluted in PBST + 1% milk in duplicate, added to plates, and incubated at room temperature for two hours. For chaotropic ELISAs, plates were next washed four times with 250 µL of 8 M urea. Plates were then incubated in anti-mouse IgG-HRP (Southern biotech) secondary antibody diluted at 1:2000 in PBST + 1% milk for one hour. The plates were then detected using 1× 3,30,5,50-Tetramethylbenzidine (TMB) (Invitrogen) and quenched with 1 M HCl after 1.5 minutes. Optical density was measured using a spectrophotometer at 450 nm and 570 nm. IC_50_ values were calculated using GraphPad Prism (GraphPad Software).

### Mice

Eight-week-old female and male C57BL/6 (strain 000664) and C3KO (strain 029661) mice were purchased from Jackson Laboratory and then bred and maintained under specific pathogen-free conditions at the University of Washington. For experiments, age and sex-matched 8-12 week old male and female mice of each genotype were used. Experiments were performed in accordance with the University of Washington Institutional Care and Use Committee guidelines (Animal Study Protocol 4283-01).

### Immunizations

Sigma adjuvant system (SAS) was mixed at a 1:1 ratio with 5 µg antigen (approximately 18 µg total nanoparticle) in PBS per manufacturer’s instructions to a final volume of 100 µL and was delivered intraperitoneally (IP). SMNP adjuvant^21^ was dosed at 5 µg per mouse formulated with 3 µg of total nanoparticle (approximately 1 µg antigen) in PBS and delivered intramuscularly (IM).

### Protein expression and tetramer generation

Recombinant full-length *Pf*CSP (SAmut-CSP) was expressed using HEK293F cells (Thermo) and then column purified using the Ni-NTA kit (Millipore Sigma). Protein was then biotinylated using the BirA500 protein ligase reaction kit (Avidity) according to the manufacturer’s instructions. Biotinylated tetramers were then generated as previously described^41^. Briefly, biotinylated proteins were incubated with streptavidin-PE at room temperature for 30 minutes. Tetramer fraction was centrifuged in a 100 kDa Amicon (Millipore) to filter out non-tetramerized protein.

### Cell isolation, enrichment, and flow cytometry

Mice were euthanized with CO_2_ asphyxiation and spleens/inguinal lymph nodes were mashed with the back of a 3 cc syringe through Nitex mesh to generate single-cell suspensions. Cells were then stained with PE-AF647 decoy reagent for 15 minutes at room temperature, followed by 30 minutes at 4°C with CSP-PE tetramer. Cells were then washed and incubated with anti-PE microbeads (Miltenyi Biotech) for 20 minutes at 4°C followed by enrichment as previously described^41^. Enriched fractions were incubated with Fc block for 15 minutes at 4°C, followed by incubation with surface markers indicated on ice for 30 minutes with antibodies listed in **Table 1**. Cells were run on a Symphony analyzer (BD) and analyzed using FlowJo software.

### ELISpot

Femurs from immunized mice were harvested and single-cell suspensions were made using a mortar and pestle filtered through Nitex mesh. Cells were then washed and resuspended in red blood cell lysis buffer (Sigma). Cells were again washed and resuspended in cell culture media (DMEM, 10% FBS, 1% pen/strep, 15 mM HEPES, 55 µM BME) and counted. 200,000 cells were then added to ELISpot plates (Millipore) that had been coated with 20 µg/mL recombinant CSP in PBS overnight. Plates were then incubated overnight at 37°C in a 5% CO_2_ incubator.

The following day (16 hours), plates were washed with PBS-T. IgG-AP antibody (Sigma) diluted 1:2000 in 1%BSA/PBS and added to each well. Plates were incubated for one hour at 37°C, washed with PBS-T, and then treated with BCIP/NBT (Mabtech) for 30 minutes at room temperature. Plates were then washed with deionized water and allowed to dry before quantification and analysis using an ImmunoSpot reader.

### Immunofluorescence

Mice were euthanized and the lymph nodes were dissected and incubated in 1% paraformaldehyde in PBS (Cytofix, BD Biosciences) overnight at 4°C. The tissue was subsequently washed twice with PBS, incubated at 4°C overnight in a 15% sucrose solution in PBS, then incubated at 4°C overnight in a 30% sucrose solution. Lymph nodes were then embedded in O.C.T. (Fisher Scientific) and frozen in a slurry of dry ice and ethanol. Following freezing, lymph nodes were cut into 10 µm sections using a cryostat (Leica Biosystems). Sections were mounted on SuperFrost Plus microscope slides (Fisher Scientific) and allowed to air dry for at least 30 minutes prior to the staining procedure. For staining, tissue sections were rehydrated in PBS for 10 minutes, fixed in 1% paraformaldehyde (Cytofix, BD Biosciences) for 5 minutes, then permeabilized in 0.5% Triton-X for 15 minutes. Sections were blocked with CAS-Block Histochemical Reagent (ThermoFisher) for at least 1 hour at 4°C in a humidified chamber. Sections were stained with antibodies in CAS-Block overnight at 4°C in a humidified chamber. Following staining, sections were washed with PBS, then mounted with Fluoromount-G (ThermoFisher). Confocal images were acquired with a Nikon C2 laser scanning confocal microscope using a 20× objective lens. Analysis of germinal center frequency and areas was performed using ImageJ.

### Parasites and mosquitoes

Transgenic *Plasmodium berghei* (strain ANKA 676m1C11, MRA-868) parasites expressing full-length (3D7 strain) *P. falciparum* CSP and a green fluorescent protein/luciferase fusion protein (Pb-PfCSP-GFP/LUC SPZ) were propagated and used to evaluate the efficacy of the PfCSP-based vaccines, as previously described^28^. Briefly, BALB/c mice were infected by intraperitoneal (IP) injection of Pb-PfCSP-GFP/LUC SPZ-infected RBCs. *Anopheles stephensi* (Nijmegen) mosquitoes were reared at the Laboratory of Malaria and Vector Research (National Institute of Allergy and Infectious Diseases, National Institutes of Health). Female mosquitoes were allowed to feed on the parasitized mice, which were anesthetized by IP injection of ketamine (50 mg/kg) and xylazine (10 mg/kg) mixture. After feeding, mice were euthanized via CO_2_ inhalation, followed by cervical dislocation. Blood-fed mosquitoes were then maintained in a humidified incubator at 19-20°C and supplied with 10% sucrose. For challenge studies, sporozoites were harvested from mosquito salivary glands at day 18-21 after an infectious blood meal, as previously described^28^.

### Immunizations and sporozoite challenge studies

RT-I53-50, RT.2-I53-50, and glycosylated RT.2-I53-50 nanoparticle immunogens were diluted in buffer to the indicated doses with SMNP adjuvants. Female B6-albino mice were immunized IM in the quadriceps at weeks 0 and 3 weeks. Challenges were conducted 3 weeks after the final immunization as indicated, where mice were challenged intravenously via the tail vein with 2,000 freshly harvested *Pb*-*Pf*CSP-GFP/LUC sporozoites as previously described^28^. Then, 40-42 hours post-challenge, mice received an IP injection of 150 µL of D-Luciferin (30 mg/mL), and were anesthetized with isoflurane (5% for induction, 1-3% for maintenance). Luciferase activity in mice was visualized through imaging of whole bodies using the IVIS® Spectrum in vivo imaging system (PerkinElmer) 10 minutes post-injection. A group of uninfected control mice were referred to as naive, and infected control mice referred to as untreated, were used for each challenge experiment. Upon completion of the experiments, mice were euthanized via CO_2_ inhalation followed by cervical dislocation. To measure the burden of parasite infection in the liver, a region of interest (ROI) in the upper abdominal area (at the location of the liver) was analyzed and the total flux or bioluminescent radiance (photons/sec) emitted by Pb-PfCSP-GFP/LUC-SPZ was calculated using the manufacturer’s software (Living Image 4.5, PerkinElmer). All animal procedures conducted at Vaccine Research Center, National Institute of Allergy and Infectious Diseases (NIAID), National Institutes of Health (NIH) were approved by the Institutional Animal Care and Use Committees of the NIAID, NIH (Animal Study Protocols VRC-23-0995 and VRC-23-1008).

### Quantitative and statistical analysis

Statistical differences between mouse groups were determined by a two-way ANOVA analysis with a Benjamini-Hochberg correction, Kruskal-Wallis test with multiple comparisons, or Mann Whitney using GraphPad Prism (GraphPad Software) as indicated.

## Acknowledgements

We are grateful to the Irvine lab for supplying SMNP for use in these studies, and to Holger Kanzler for constructive discussions. This research was supported by National Institutes of Health grant F32AI178963 (C.E.M) and the Gates Foundation (INV-043758 to M.P. and N.P.K.). This study was funded, in part, by the Intramural Research Program of the National Institutes of Health (NIH). The contributions of the NIH authors are considered works of the United States Government. The findings and conclusions presented in this paper are those of the authors and do not necessarily reflect the views of the NIH or the U.S. Department of Health and Human Services. Flow cytometry data was acquired through the University of Washington Cell Analysis Facility Shared Resource Lab with NIH award 1S10OD024979-01A1 funding for the Symphony A3.

## Competing Interests

The authors declare no competing interests.

## Notes

### Competing Interest Statement

The authors have declared no competing interest.

