## Supplemental Data for "Mannosylated nanoparticle immunogens enhance the circumsporozoite protein-specific B cell response and improve protection against sporozoite challenge"

### Supplementary information

**Supplementary Table 1. Amino acid sequences of novel proteins used in this work.**

| |
|---|
| >I53-50A-4gly<br>MDSKGSSQKGSRLLLLLVVSNLLLPQGVLAKIEELFKKHKIVAVLRANSVEEAIEKAVAVFAGGVHLIEITF<br>TVPNATTVIKALSVLKEKGAIIGAGTVTSVEYANETVESGAEFIVSPHLDEEISNFTKEKGVFYMPGVMTP<br>TELVKAMKLGHTILKLFPGEVVGPQFVKAMKGPFHNATFVPTGGVNLDNVCEWFKAGVLAVGVGSALV<br>KGTPDEVREKAKAFVEKIRGCTEGSHHHHHH |
| >I53-50A-3gly<br>MDSKGSSQKGSRLLLLLVVSNLLLPQGVLAKIEELFKKHKIVAVLRANSVEEAIEKAVAVFAGGVHLIEITF<br>TVPDADTVIKALSVLKEKGAIIGAGTVTSVEYANETVESGAEFIVSPHLDEEISNFTKEKGVFYMPGVMT<br>PTELVKAMKLGHTILKLFPGEVVGPQFVKAMKGPFHNATFVPTGGVNLDNVCEWFKAGVLAVGVGSAL<br>VKGTPDEVREKAKAFVEKIRGCTEGSHHHHHH |
| >RT.2-I53-50A-4gly<br>MDSKGSSQKGSRLLLLLVVSNLLLPQGVLASSNSKMDPNANPNANPNANPNANPNANPNANPNANPN<br>ANPNANPNANPNANPNANPNANPNANPNANPNANPNANPNANPNKNNQGNGQGHNMPNDPNRNV<br>DENANANSAVKNNNNEEPSDKHIKEYLNKIQNSLSTEWSPCSVTCGNGIQVRIKPGSANKPKDELDYA<br>NDIEKKICKMEKCSSVGGSGGSGSEKAAKAEEAARKIEELFKKHKIVAVLRANSVEEAIEKAVAVFAGGV<br>HLIEITFTVPNATTVIKALSVLKEKGAIIGAGTVTSVEYANETVESGAEFIVSPHLDEEISNFTKEKGVFYMP<br>GVMTPTELVKAMKLGHTILKLFPGEVVGPQFVKAMKGPFHNATFVPTGGVNLDNVCEWFKAGVLAVG<br>VGSALVKGTPDEVREKAKAFVEKIRGCTEGSHHHHHH |
| >RT.2-I53-50A-3gly<br>MDSKGSSQKGSRLLLLLVVSNLLLPQGVLASSNSKMDPNANPNANPNANPNANPNANPNANPNANPN<br>ANPNANPNANPNANPNANPNANPNANPNANPNANPNANPNANPNKNNQGNGQGHNMPNDPNRNV<br>DENANANSAVKNNNNEEPSDKHIKEYLNKIQNSLSTEWSPCSVTCGNGIQVRIKPGSANKPKDELDYA<br>NDIEKKICKMEKCSSVGGSGGSGSEKAAKAEEAARKIEELFKKHKIVAVLRANSVEEAIEKAVAVFAGGV<br>HLIEITFTVPDADTVIKALSVLKEKGAIIGAGTVTSVEYANETVESGAEFIVSPHLDEEISNFTKEKGVFYM<br>PGVMTPTELVKAMKLGHTILKLFPGEVVGPQFVKAMKGPFHNATFVPTGGVNLDNVCEWFKAGVLAV<br>GVGSALVKGTPDEVREKAKAFVEKIRGCTEGSHHHHHH |

**Supplementary Table 2. Flow cytometry reagents used in this work.**

| Antibody | Clone | Color | Source |
|---|---|---|---|
| anti-mouse B220 | RA3-6B2 | BUV737 | BD Biosciences |
| anti-mouse CD4 | GK1.5 | BUV805 | BD Biosciences |
| anti-mouse CD138 | 281-1 | BV650 | BD Biosciences |
| anti-mouse CD38 | 90 | AF700 | Invitrogen |
| anti-mouse GL7 | GL7 | eF450 | Invitrogen |
| anti-mouse IgM | II/41 | BV786 | BD Biosciences |
| anti-mouse IgD | 11-26c.2a | BUV395 | Biolegend |
| anti-mouse CD73 | eBioTY/11.8 | PE-Cy7 | Invitrogen |
| anti-mouse CD80 | 16-10A1 | BV605 | BD Biosciences |
| anti-mouse Fc | 2.4G2 | | BD Biosciences |

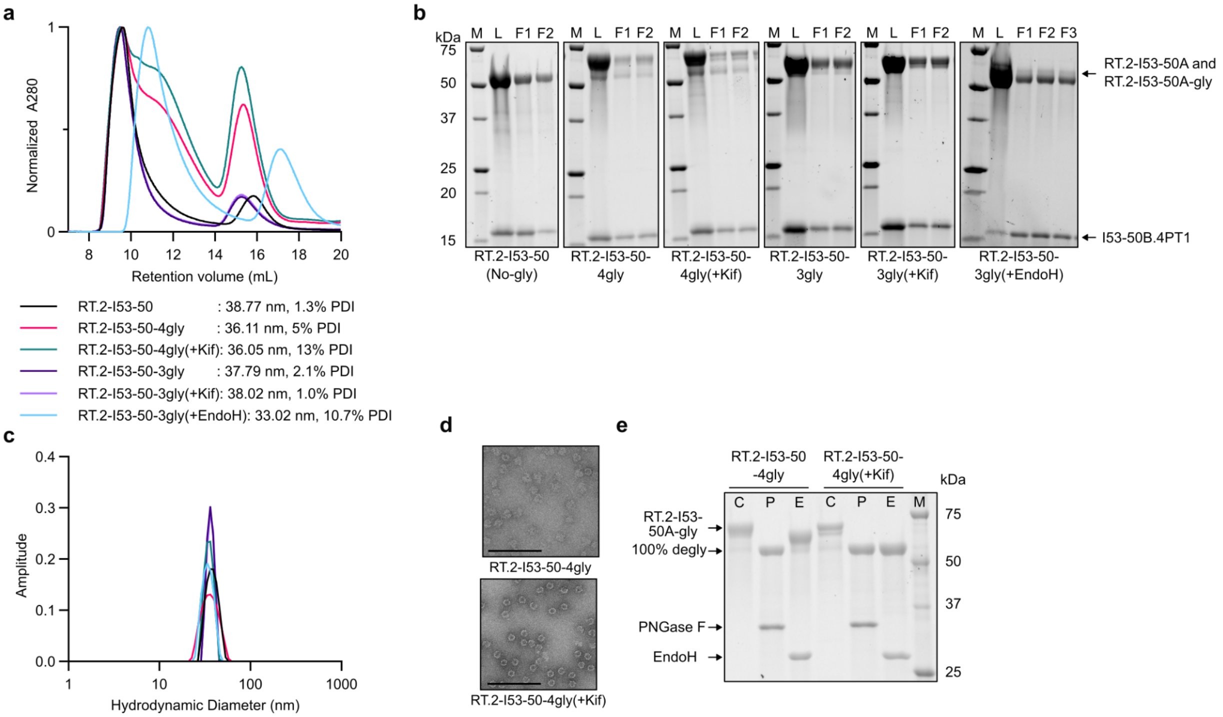


**Supplementary Figure 1. Characterization of RT.2-I53-50 nanoparticles and glycoprofiling of RT.2-I53-50A-4gly. a** Size exclusion chromatography (SEC) chromatograms on a Superose 6 Increase 10/300 GL column. The first peak corresponds to the assembly, and the second peak corresponds to excess components. RT.2-I53-50A-3gly(+EndoH) trace is shifted rightward by 2 mL due to a different ÄKTA FPLC protocol injection time. **b** SDS-PAGE for bare and glycosylated RT.2-I53-50 nanoparticles; M = marker, L = load, F1 & F2 = fractions pooled for immunization studies. **c** Hydrodynamic diameter (nm) of nanoparticles as determined by dynamic light scattering (DLS). **d** Negative-stain electron micrographs for RT.2-I53-50-4gly nanoparticles at 57k magnification. Black bar inset in the lower left for each micrograph corresponds to 200 nm size. **e** Control untreated (C), PNGase F (P), and Endo H (E) treated glycosylated RT.2-I53-50-4gly nanoparticles. SDS-PAGE was run under denaturing conditions.

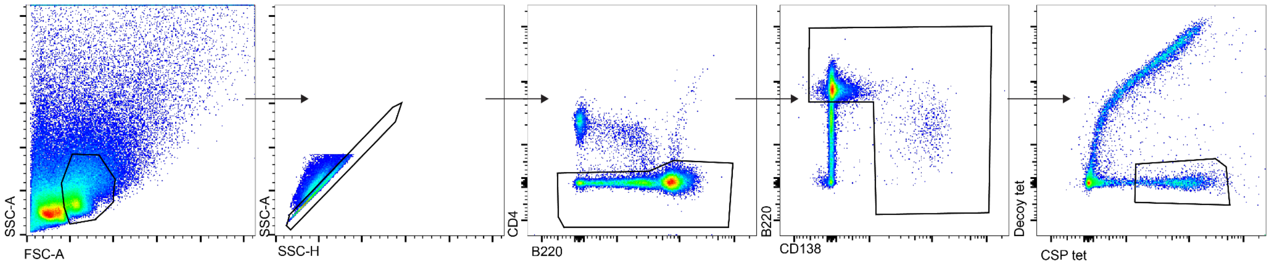


**Supplementary Figure 2. Gating strategy.** Shown is the gating strategy to identify CSP+ B cells in C57BL/6 mice immunized with 3gly-mannose(+Kif) 8 days prior. Gated on live lymphocytes, singlets, CD4-, B cells (B220+ or CD138+), CSP tetramer+.

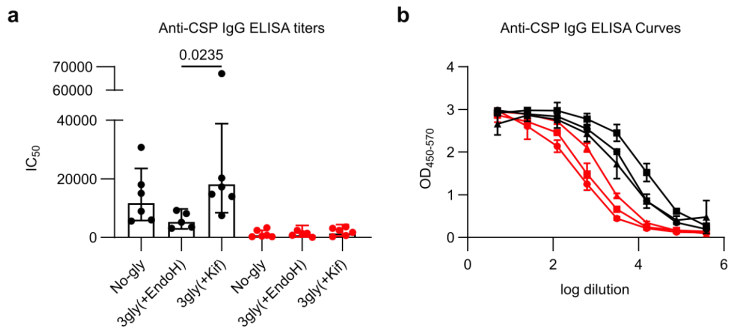


**Supplementary Figure 3. Glycosylation does not influence early high-affinity antibody titers.** C57BL/6 or C3-deficient (C3KO) mice were immunized intramuscularly with 3 μg of No-gly, 3gly(+EndoH), or 3gly(+Kif) nanoparticles adjuvanted with SMNP adjuvant. Eight days later, blood was harvested for analysis. **a** Serum CSP-specific IgG titers and **b** curves quantified by ELISA. Statistical analysis by 2-way ANOVA with a Benjamini-Hochberg correction. Mean and standard deviation are plotted.

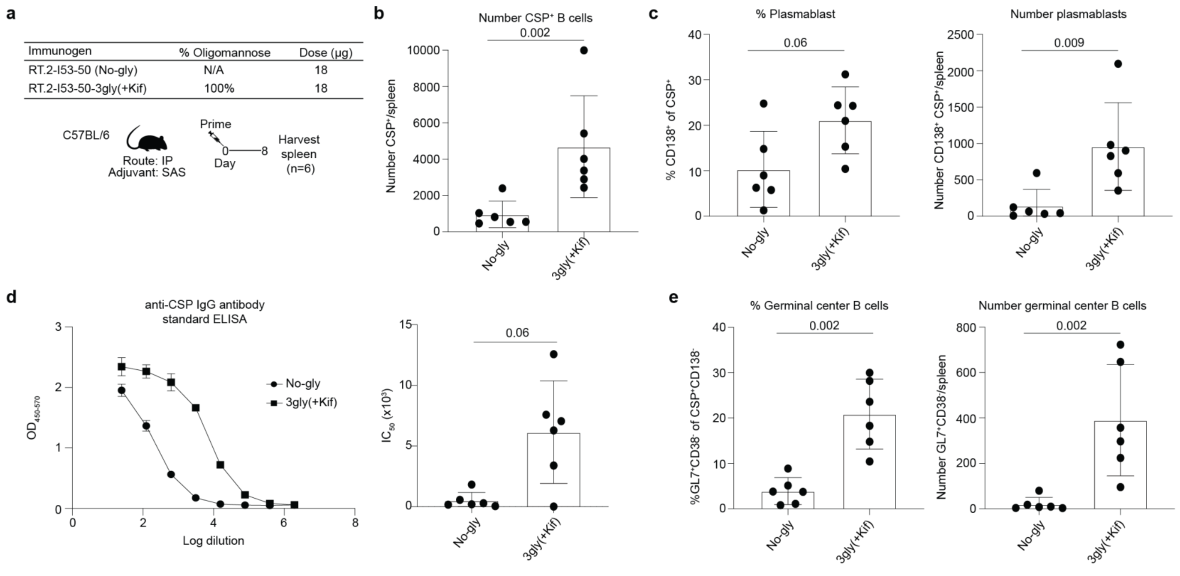


**Supplementary Figure 4. Mannosylated NPs improve CSP-specific responses across route, dose, and adjuvant. a** C57BL/6 mice were immunized intraperitoneally with 18 µg of No-gly or 3gly(+Kif) nanoparticles with SAS adjuvant. 8 days later spleens were harvested. **b** The number of $CSP^+$ B cells. **c** The proportion and number of CSP-specific PBs ($CSP^+CD138^+$). **d** Serum CSP-specific IgG was quantified by ELISA. **e** The proportion and total number of CSP-specific GC B cells ($CSP^+CD138^-CD38^-GL7^+$). n=6. Statistical analysis by Mann-Whitney.
